# Identification of donor landraces for early vegetative stage salinity tolerance in wheat by employing multivariate selection indices

**DOI:** 10.1101/2025.09.10.675370

**Authors:** Jagadhesan B, Arvind Kumar, Shailendra K Jha, Ramesh R, Rakesh Pandey, Sanjay Kalia, Renu Pandey, Amit Singh, Sundeep Kumar, Gyanendra P Singh, Viswanathan Chinnusamy, Lekshmy Sathee

## Abstract

Soil salinity (EC of ≥4 dS m^-1^) is a major abiotic stress leading to decreased crop production due to osmotic, ionic, and cellular ionic imbalance balance. The current study aims to identify salinity-tolerant wheat landraces by investigating morpho-physiological traits. Three hundred fifty wheat landraces and check lines were evaluated for salt stress (15 dS m^-1^) tolerance in a hydroponic set-up. The clustering analysis categorized 350 genotypes into four control and 15 dS m^-1^ conditions clusters. Across the multi-trait genotype-ideotype distance index (MGIDI), Smith Hazel Index (SHI), and Stress Susceptibility Index (SSI), seven genotypes were consistently identified as tolerant among the top 150. In comparison, 12 genotypes were consistently classified as least tolerant. These identified tolerant landraces serves as important resources and novel salt tolerance donors for wheat. The tolerant*sensitive, tolerant* tolerant check, crosses can be used for generating RIL populations to delineate genetic basis of salt tolerance of newly identified lines.

## 1. Introduction

Among the many abiotic stresses crops come across due to the sedentary habit is soil salinity which dampens the productivity incrementally each year. Soil salinity is pressing issue for about 6% of the world’s land area, which includes 20% of arable land and 33% of irrigated land (Safdar *et al*., 2019). Every year, 10 million ha of farmland is destroyed by salt buildup caused by human activities and other climate change-related factors (Isayenkov and Maathuis, 2019). Natural salt stress has a devastating effect on the development and yield of sessile plants (Alam *et al*., 2019). Excess salts in the rhizosphere hinder water and nutrient absorption. Salinity stress decreases seed germination, plant growth, yield and manifests as changes in physiological and biochemical responses (Ahanger *et al*., 2019). The majority of field crops, such as rice *(Oryza sativa)*, wheat *(Triticum aestivum)*, maize *(Zea mays)*, and others, are classified as glycophytic plants which are sensitive to soil salinity. Nevertheless, the techniques for adapting to salt stress are the same regardless of whether the plants in question are glycophytes or halophytes (Yokoi *et al*., 2002). Wheat is a very significant agricultural plant on a global scale and provides sustenance for a substantial population. Nevertheless, wheat exhibits only a low level of tolerance to salt, resulting in a reduction of over 60% in its grain production in saline conditions (Khan *et al*., 2016). To effectively address this problem, one of the most successful solutions is the development of salt-tolerant cultivars (Oyiga *et al*., 2016). The identification of genotypes that have a broad range of adaptability to salinity is a minimum need. Despite this, only a small number of wheat genotypes that have been discovered as being salt-tolerant have been extensively exploited in breeding for salt-tolerant cultivars (Chatrath *et al*., 2007). Therefore, it is necessary to discover novel germplasms that are tolerant to salt to extend the gene base and to give elite resources that have genetic backgrounds that are extensively adaptable.

The salt tolerant processes in plants have intricate characteristics. Initially, salt will impact the plants’ performance via osmotic and ionic stress, and finally with oxidative stress (Zeng *et al*., 2015). Plants increase their ability to withstand osmotic stress by preserving cell turgor via the buildup of soluble compounds that aid in osmotic adjustments, such as proline, glycine betaine, and soluble sugars (de Freitas *et al*., 2019). Concurrently, the ability of tissues to withstand the harmful effects of ionic toxicity may be enhanced by controlling the balance of sodium and potassium ions inside cells and by isolating and preventing the accumulation of sodium ions (Arif *et al*., 2020). Furthermore, the process involves the production of anti-oxidant enzymes and metabolites that eliminate reactive oxygen species (ROS) in response to oxidative stress (Gapińska *et al*., 2008).

A lack of efficient assessment methodologies has hindered the screening of wheat genotypes for salt tolerance (Li *et al*., 2020). Biomass yield is a valuable indication since it allows for directly assessing economic gain in the presence of salt stress (Munns and James, 2003). Furthermore, it is important to note that the ability of plants to tolerate salt is dependent on the particular stage of development, and the sensitivity of physiological characteristics to salt changes throughout various growth stages. This complicates the process of choosing salt-tolerant indicators and results in a decrease in selection efficiency (Zeng *et al*., 2003). Although the overall agronomic criteria have a poor selection efficiency, multiple studies have successfully evaluated salt tolerance in wheat using multivariate analysis (El-Hendawy *et al*., 2005; Hasan *et al*., 2015). Plant breeding primarily focuses on agronomic traits, with a particular emphasis on maximizing output in agricultural production.

Considering all the facts mentioned above, we believe that combining biomass ranking with other salt stress-associated traits and integrated ranking would aid in the efficient assessment of salt-tolerant genotypes. This study aimed to analyze the salt tolerance of 350 wheat genotypes collected from different regions of the country during the seedling stage. The objective was to find salt-tolerant genotypes using MGIDI, SH, and SSI. An assessment was conducted on wheat genotypes to examine the impact of salt stress on important physiological characteristics such as leaf chlorophyll content, cell membrane stability. The identified genotypes are potential candidates for production and may contribute to a deeper knowledge of the processes underlying salt tolerance in wheat.

## 2. Materials and methods

### 2.1 Plant material and Experimental setup and design

A diversity based mini core collection of wheat landraces contains 350 accessions including check lines like Kharchia-65 (K-65), KRL-210, HD3226, and HD2851 seeds were procured from Borlaug Institute for Southern Asia (BISA) (India) and used for this experiment **(Supplementary Table 1.).** The seeds were subjected to seed treatment and grown in a salt solution for thirty days to, measure the agro-physiological traits following the protocol standardized by us earlier (Boopal *et al*., 2023) (Supplementary.

Experiment was conducted in a glass house chamber under the controlled conditions of the National Phytotron Facility, IARI, New Delhi. Seedlings were grown with 15 dSm^-1^ treatment comprising of combination of three salts: NaCl, CaCl_2_.2H_2_O, and Na_2_SO_4_ along with control. The seeds of wheat genotype were subjected to surface sterilization using sodium hypochlorite (NaClO, 2%), and any residual traces were eliminated by rinsing with deionized water four to five times. Later, the surface sterilized seeds were enveloped in moistened cellulosic germination paper. After 7 days healthy seedlings with similar growth were selected for transplanting. The seedlings were held in acrylic sheets with foam support and placed in plastic trays filled with 20 liters of modified Hoagland nutrient solution. Each hill contained two seedlings, and there were three replication trays for each treatment. Every 3-4 days, the Hoagland nutrient solution was replaced, and the pH was maintained between 5.2 to 5.6 using 0.1 M HCl/ 0.1 M NaOH throughout the study and a pocket pH meter was used to record the pH of the nutrient solution. The modified Hoagland nutrient solution was prepared in de-ionized RO water. The concentration of salts and the EC were adjusted using a handheld EC meter. In the Wheat Glasshouse facility, the temperature was maintained at 22 °C during the day to 12 °C at night. The photoperiod lasts for 10 hours during the day and 12 hours during the night, with a photon flux density of 400 μmol m^−2^ s^−1^. The relative humidity in the glasshouse varies from 80% to 90%. Seedlings were allowed to grow for 30 days in hydroponics with recommended changes in modified Hoagland nutrient solution and treatments as described above.

### 2.2 Plant height and fresh and dry biomass

Plant height of each genotype was measured with a standard ruler from every replicated tray from thirty days of old seedlings. Fresh and dry biomass of the seedlings were measured using weighing scale at the time of harvest and after achieving constant dry weight by oven drying respectively.

### 2.3 Green leaf area

The green leaf area of thirty-day-old seedlings was measured by using a Leaf Area Meter LiCOR 3100 model manufactured by Lincon in Nebraska, USA.

### 2.4 Membrane stability index

Thirty-day-old seedlings were harvested, and the membrane stability index (MSI) was computed and calculated following the earlier procedure (Sairam *et al*., 1997).

### 2.5 Chlorophyll content index and Relative water content

A non-destructive mode chlorophyll content was measured in thirty-day-old seedlings using Chlorophyll content meter (Boopal *et al*., 2023) and relative water content (RWC) and water deficit (WD) was measured as described by (Weatherley, 1950).

### 2.6 Chlorophyll fluorescence

The seedlings’ chlorophyll fluorescence of the leaves was assessed using a handheld chlorophyll fluorometer (Model: OS5p+ Advance Chlorophyll Fluorometer). To establish a dark-adapted condition, a clip was placed in the center of the upper most expanded leaf for 20 minutes. Photosystem II (PSII) efficiency was then determined by calculating the Fv/Fm ratio and Fv/F0, following the methodology described by (Krause and Weis, 1991; Mohammed *et al*., 1995).

### 2.7 Thermal imaging and canopy temperature

The canopy temperature (CT) of thirty days old seedlings was recorded by using via an infrared thermal camera (Fluke TiX660 model, Fluke Infrared camera, Everett, Washington, USA (Boopal *et al*., 2023) and images were analyzed by using software package SmartView 4.3TM Researcher Pro (Fluke Thermography, Plymouth, MN, USA) and exported to MS Word for further analysis (Naguib *et al*., 2021).

### 2.8 Statistical analysis

Homogeneity of data was performed using the R program. R program was also used to compute a two-way analysis of variance (ANOVA) with genotype, control, and salt treatment effects to compute adjusted P values and significance levels. Graphs were made using the R program, and ‘GraphPad Prism version 9.0.0’ (La Jolla, California, USA). A correlation plot was created using the ‘corrplot’ package in the R program. Principle component analysis (PCA) was performed using the ‘FactoMineR’ and ‘factoextra’ packages in the R program. The hierarchical cluster analysis procedure was utilized to group cases or variables with similar characteristics in a dataset by using ‘hclust’ R package. Multi-trait genotype ideotype distance index (MGIDI) to assess and identify the most superior genotypes, considering their quantitative characteristics, and this was performed by the functions ‘gamam’ and ‘mgidi’ of the package ‘metan’ of the R program were used for the MGIDI and calculation (Olivoto and Nardino, 2021). Smith-Hazel Index was computed by the “metan” package in the R program. Stress Susceptibility Index (SSI) was computed according to the formula given by (Fischer and Maurer, 1978). Venn diagram was constructed using the Bioinformatics & Evolutionary Genomics online tool.

## 3. Results

### Assessment of morphophysiological parameters associated with early vegetative stage salinity tolerance of wheat landrace minicore

The effect of salinity stress (15 dSm^-1^) on shoot length (SL), root length (RL), and plant height (PH) is given in **(Fig. 1 (a, b, c)).** The salt treatment significantly affected SL, RL, and PH of 350 wheat landraces, including tolerant check (K-65). A significant decrease in SL, RL, and PH was observed under 15 dSm^-1^ compared to the control. In 15 dSm^-1^, SL ranged from 19.1 cm (4687) to 56.1 cm (IC532495) with a mean of 34.60 cm, while RL ranged between 10.23 cm (IC75217) and 36.23 cm (IC321153) with a mean of 18.82cm. PH ranged from 33.60 cm (4379) to 82.83 cm (IC532495), with a mean of 53.42 cm in 15 dSm^-1^. The calculated F-values for SL, RL, and PH were 4.08, 5.52, and 4.08, respectively. The effect of salinity stress (15 dSm^-1^) on CCI, RWC, and WD is given in **(Fig. 1 (d, e, and f))**. Salt treatment significantly affected CCI, RWC, and WD of 350 wheat landraces, including check (K-65). A significant reduction in CCI, RWC, and WD was seen under 15 dSm^-1^ treatment concerning the control, with corresponding F-values of 3.43, 13.02, and 13.05. In 15 dSm^-1^, CCI ranged from 0.00 (IC551375) to 4.33 (IC111914) with a mean of 1.98, while RWC ranged between 27.75 % (IC531275) and 91.49 % (IC418402) with a mean of 69.71 %. WD ranged from 8.50 % (IC418402) to 72.24 % (IC531275), with a mean of 30.2355 % in 15 dSm^-1^.

**Fig 1.**
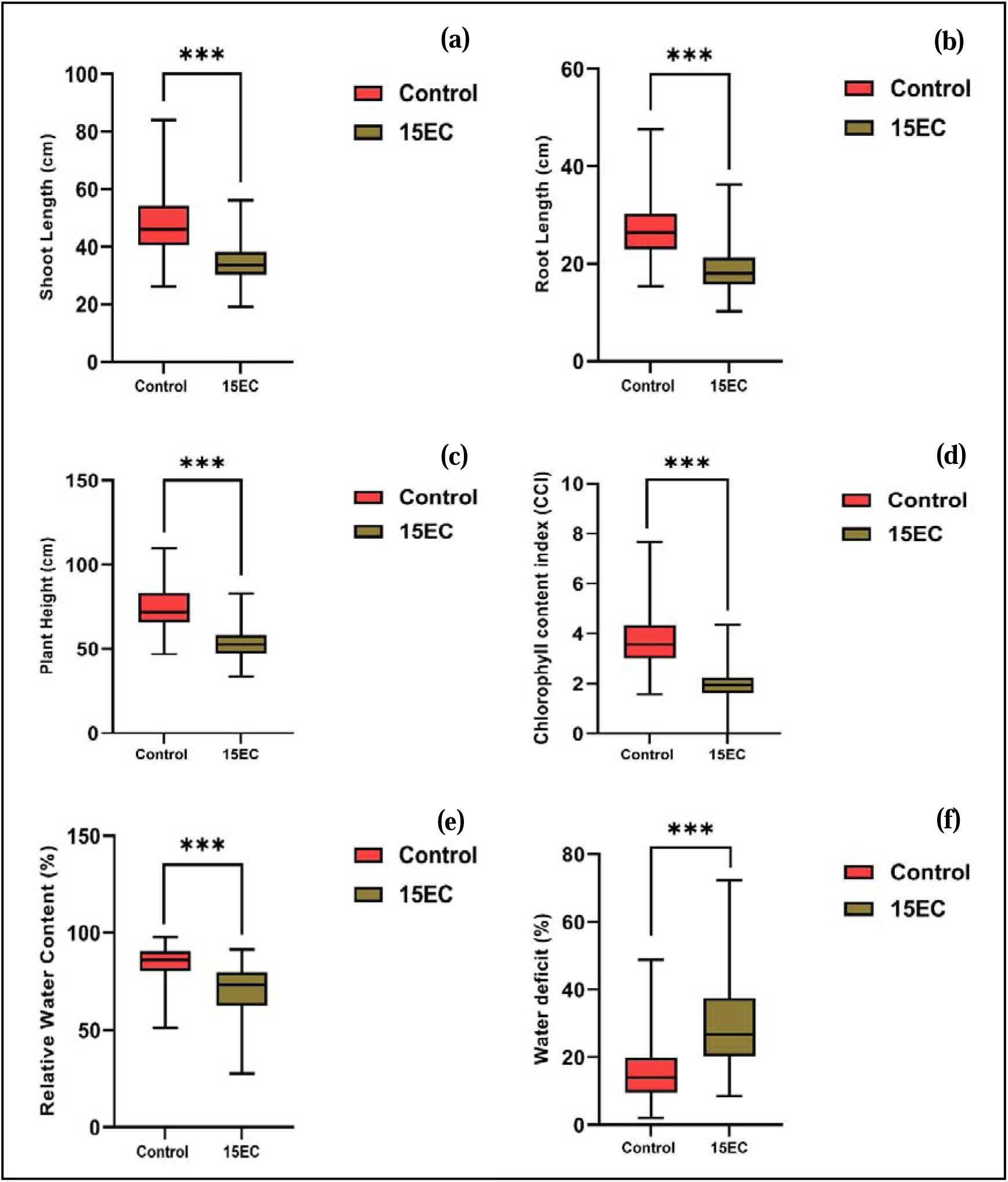
Effect of salinity stress (Control, and 15 dSm^-1^) on shoot length (a), root length (b), plant height (c) chlorophyll content index (d) relative water content (e) and water deficit (f) of 350 wheat genotypes including check (Kharchia 65). Salinity stress was imposed for 4 weeks as mix of salts to 7 days old wheat seedlings grown under hydroponics. Asterisk *, ** and *** indicate 5%, 1% and 0.1% levels of significance, respectively.

The effect of salinity stress (15 dSm^-1^) on shoot fresh weight (SFW), root fresh weight (RFW), and total fresh weight (TFW) is given in **(Fig. 2 (a, b, c))**. The salt treatment significantly affected the 350 landraces, including check. A significant reduction in SFW, RFW, and TFW was seen under 15 dSm^-1^ compared to the control, with corresponding F-values of 4.70, 6.16, and 5.33. In 15 dSm^-1^, SFW ranged from 0.17 g (IC582907) to 2.46 g (IC335977) with a mean of 0.79 g, while RFW ranged between 0.07 g (4379) and 1.06 g (IC598261) with a mean of 0.30 g. TFW ranged from 0.30 g (4379) to 3.14 g (K-65), with a mean of 1.10 g in 15 dSm^-1^. The effect of salinity stress on shoot dry weight (SDW), root dry weight (RDW), and total dry weight (TDW) is given in **(Fig. 2 (d, e, f))**. As with other measured traits, under salinity, a significant reduction of dry biomass was recorded to control. A significant reduction in SDW, RDW, and TDW was seen under 15 dSm^-1^ compared to the control group, with corresponding F-values of 4.64, 7.78, and 5.17. In 15 dSm^-1^, SDW ranged from 0.042 g (IC589296) to 0.49 g (IC335977) with a mean of 0.16 g, while RDW ranged between 0.007 g (IC29002) and 0.093g (K-65) with a mean of 0.03 g. TDW ranged from 0.06 g (IC75217) to 0.574 g (IC335977), with a mean of 0.194 g in 15 dSm^-1^.

**Fig 2.**
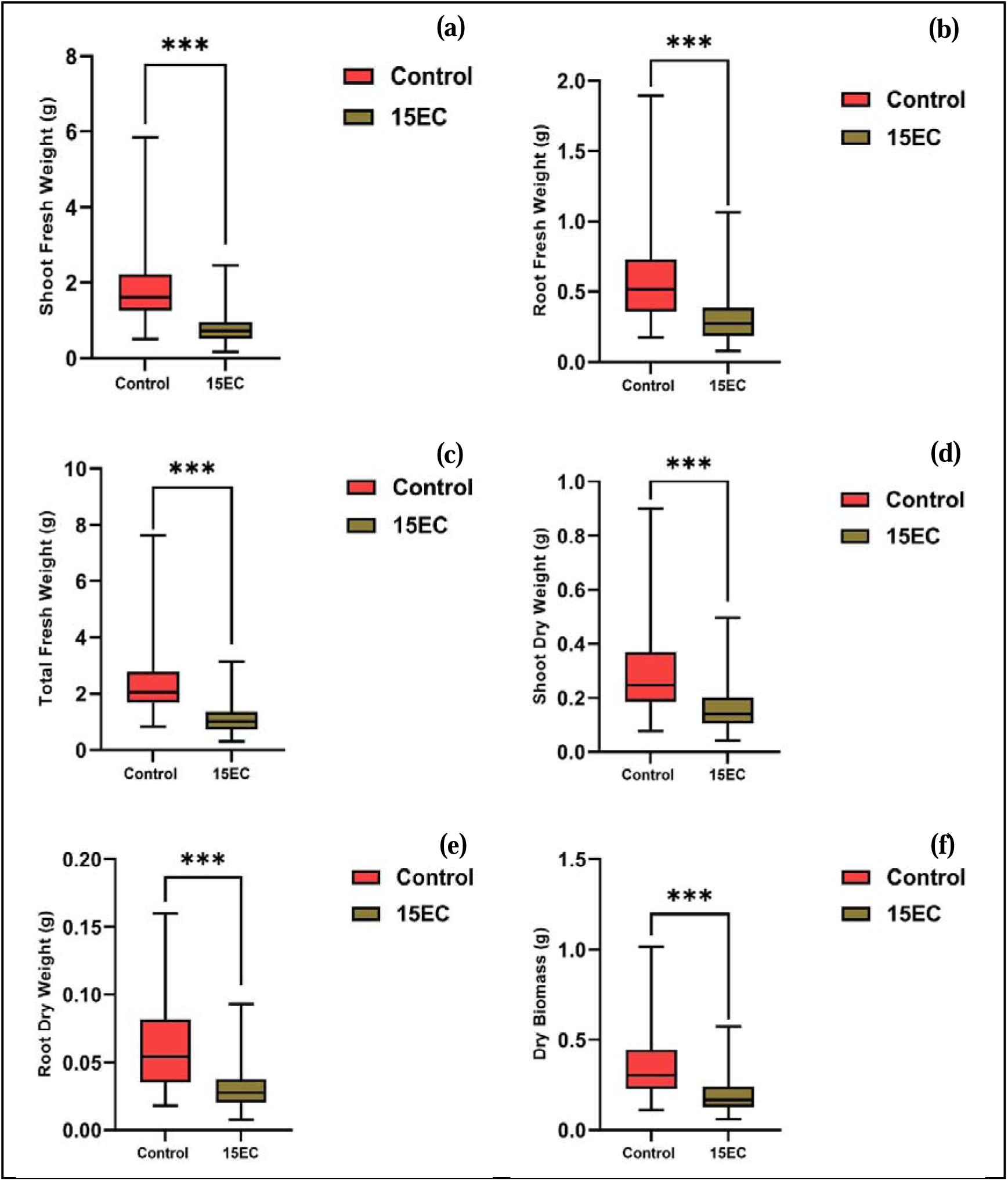
Effect of salinity stress (Control, and 15 dSm^-1^) on shoot fresh weight (a), root fresh weight (b), total fresh weight (c) shoot dry weight (d) root dry weight (e) and total dry weight (f) of 350 wheat genotypes including check (Kharchia 65). Salinity stress was imposed for 4 weeks as mix of salts to 7 days old wheat seedlings grown under hydroponics. Asterisk *, ** and *** indicate 5%, 1% and 0.1% levels of significance, respectively.

The effect of salinity stress (15 dSm^-1^) on green leaf area (GLA), visual scoring (VS), and membrane stability index (MSI) is given in **(Fig. 3 (a, b, c)).** A significant reduction in GLA, VS, and MSI was seen under 15 dSm^-1^ compared to the control, with corresponding F-values of 11.14, 37.39, and 8.21. In salinity treatment, GLA ranged from 0.00 cm^2^ (no: 27 genotypes; No green leaves were recorded) to 44.25 cm^2^ (K-65) with a mean of 12.82 cm^2^, while VS ranged between 3 (IC598266) and 9 (IC145983) with a mean of 6. MSI ranged from 5.95 % (IC7145972) to 71.06 % (IC585562), with a mean of 29.92 % in 15 dSm^-1^. The effect of salinity stress (15 dSm^-1^) on Canopy temperature (CT), Fv/Fm, and Fv/F0 is given in **(Fig. 3 (d, e, and f)).** The salt treatment significantly affected CT, Fv/Fm/F0 of 350 wheat landraces, including check (K-65). A significant reduction in Fv/Fm and Fv/F0 was seen under 15 dSm^-1^ compared to the control group, with corresponding F-values of 4.38, 57.0, and 6.56. The CT ranged from 18.53 °C of (IC443767) genotypes to 32.76 °C (IC0290196) with a mean of 26.83 °C, while Fv/Fm ranged between 0 (IC397815) and 0.827 (IC421880) with a mean of 0.699 in 15 dSm^-1^. In the case of Fv/F0, ranged from 0 (IC397815) to 5.017 (IC317610) with a mean of 3.254 in 15 dSm^-1^.

**Fig 3.**
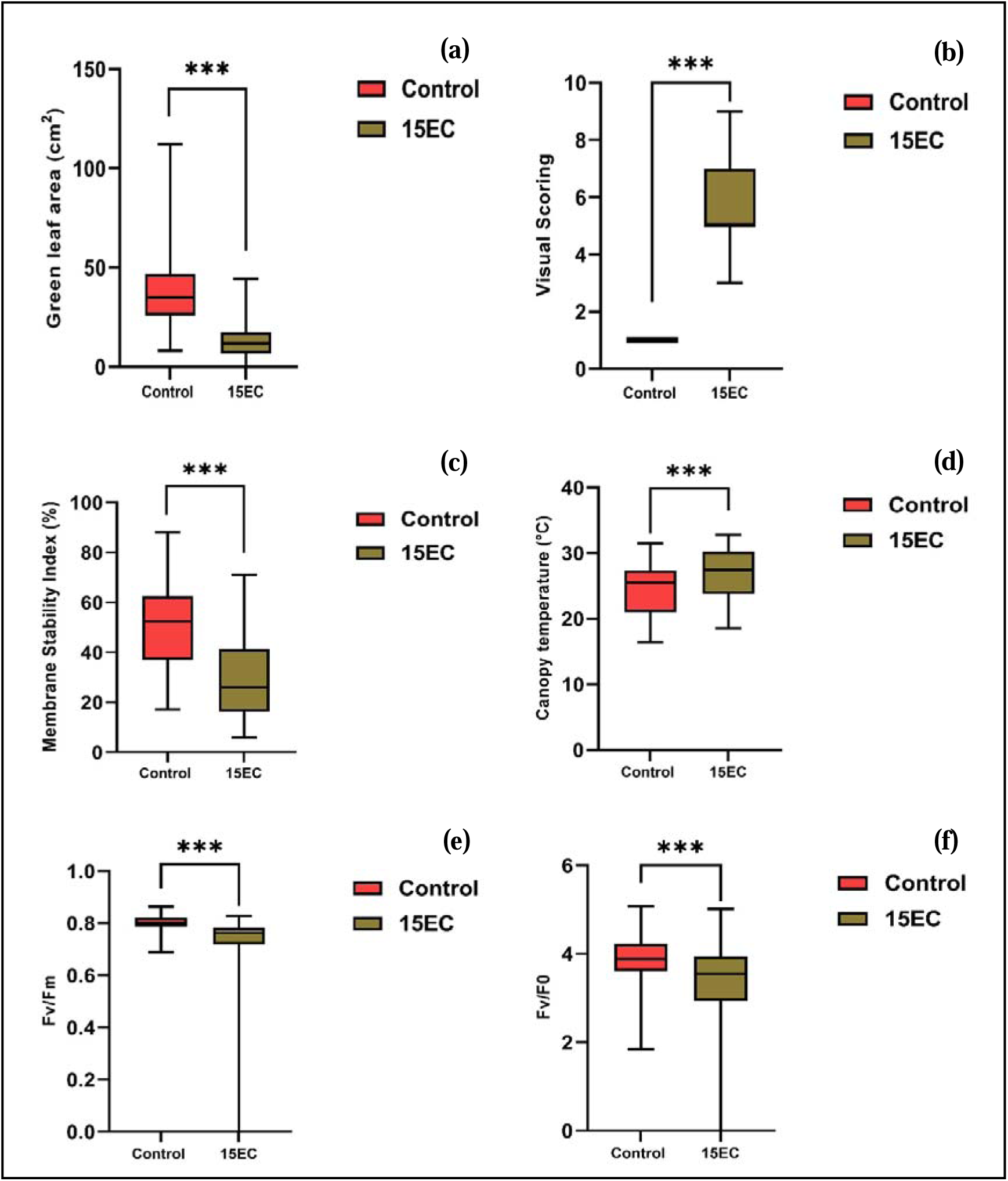
Effect of salinity stress (Control, and 15 dSm^-1^) on green leaf area (a), visual scoring (b), membrane stability index (c) canopy temperature (d) Fv/Fm (e) and Fv/F0 (f) of 350 wheat genotypes including check (Kharchia 65). Salinity stress was imposed for 4 weeks as mix of salts to 7 days old wheat seedlings grown under hydroponics. Asterisk *, ** and *** indicate 5%, 1% and 0.1% levels of significance, respectively.

### Pearson correlation of salinity tolerance associated traits

Correlation among the traits of thirty-day-old seedlings grown in control and 15 dSm^-1^ was determined **(Fig. 4 (a, b)).** In control, most of the traits were positively correlated **(Fig. 4 (a)).** Dry biomass (TDW) is highly positively correlated with SDW (r = 0.99), TFW (r = 0.82), GLA (r = 0.71), SFW (r = 0.79), RL (r = 0.38), PH (r = 0.74), SL (r = 0.72), CCI (r = 0.36), WD (r = 0.31), CT (r = 0.37), MSI (r = 0.37), Fv/Fm (r = 0.33), and Fv/F0 (r = 0.35), and negatively correlated with RWC (r = -0.31). RWC was negatively correlated with most of the traits, SDW (r = -0.34), TFW (r = -0.31), GLA (r = -0.28), SFW (r = -0.29), RL (r = -0.26), PH (r = -0.35), SL (r = -0.30), CCI (r = -0.10), WD (r = -1.00), CT (r = -0.18), MSI (r = -0.11), Fv/Fm (r = -0.31), and Fv/F0 (r = -0.27). In 15 dSm^-1^ conditions, many traits were positively correlated **(Fig. 4 (b)).** Dry biomass (TDW) is highly positively correlated with SDW (r = 1.00), TFW (r = 0.92), GLA (r = 0.61), SFW (r = 0.91), RL (r = 0.52), PH (r = 0.78), SL (r = 0.75), CCI (r = 0.16), WD (r = 0.25), CT (r = 0.29), MSI (r = 0.19), Fv/Fm (r = 0.29), and Fv/F0 (r = 0.32), and negatively correlated with RWC (r = -0.25) and VS (r = - 0.32). RWC was negatively correlated with most of the traits, SDW (r = -0.26), TFW (r = -0.23), SFW (r = -0.22), RL (r = -0.28), PH (r = -0.30), SL (r = -0.23), WD (r = -1.00), CT (r = -0.24), Fv/Fm (r = -0.03), and Fv/F0 (r = -0.16), and positively correlated with GLA (r = 0.03), CCI (r = 0.16), MSI (r = 0.18). K-means cluster analysis was performed to do a clustering of all the genotypes grown in control and salinity **(Supplementary Fig. 1 (a, b))**. Genotypes are clustered based on the relatedness in the mean value of the traits. Here, the K value is fixed into 4; the results will be divided into 4 clusters. Four individual clusters of 350 genotypes are produced under controlled conditions and 15 dSm^-1^. In control, clusters I, II, III, and IV contain 31, 57, 146, and 104 genotypes, respectively **(Supplementary Fig. 1 (a)).** When a more significant number of genotypes are clustered in group III, it leads to a higher similarity in the mean values of measured attributes among these genotypes. In 15 dSm^-1^, clusters I, II, III, and IV viz contain 14, 83, 25, and 216 genotypes, respectively **(Supplementary Fig. 1 (b)).** While a larger number of genotypes are grouped in cluster IV, it increases similarity in the average values of the measured characteristics among these genotypes.

**Fig 4.**
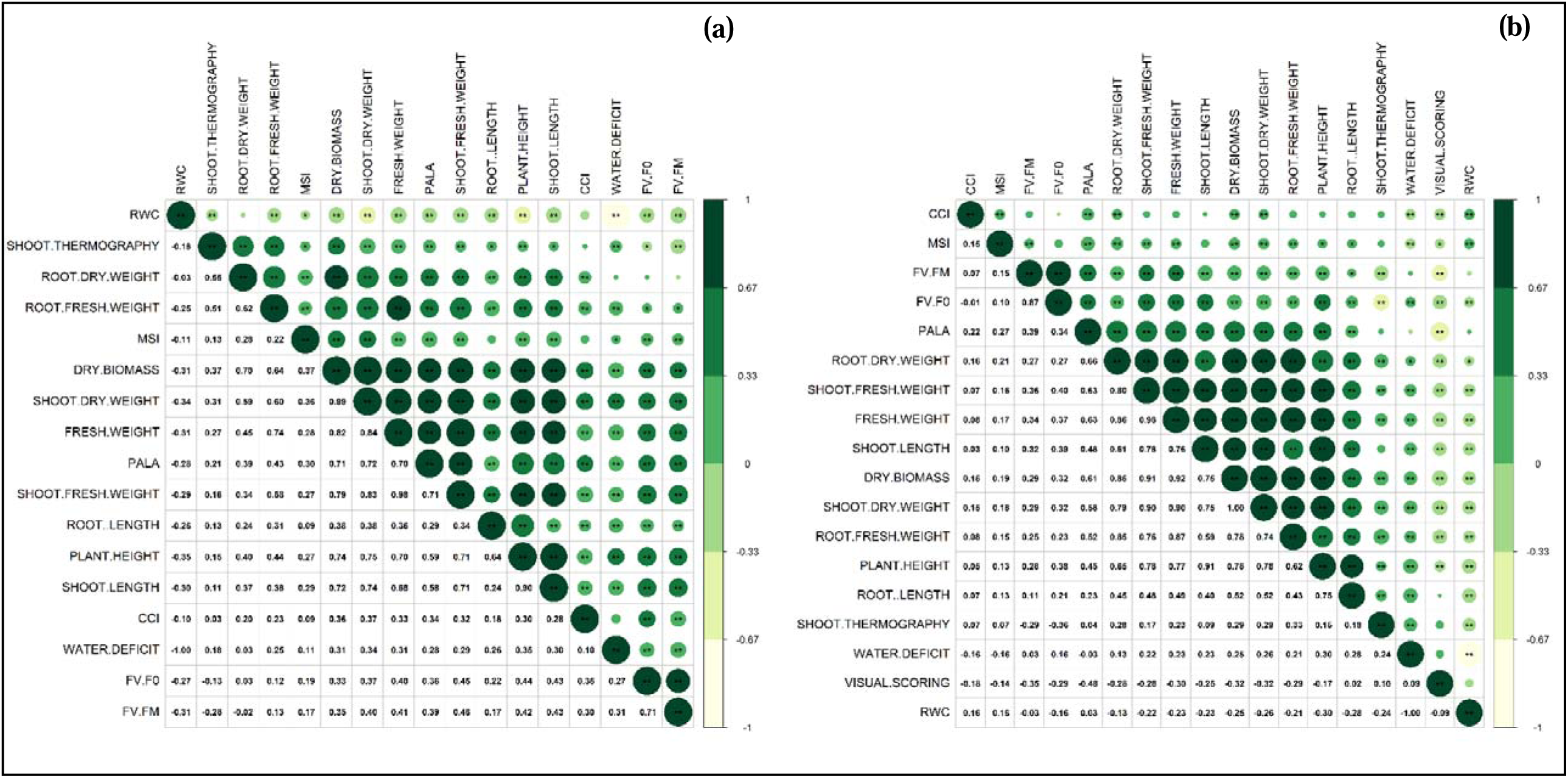
Corrplot representing the Pearson correlation matrix showing correlation between traits recorded under salinity stress. Control (a) and salt treatment 15 dSm^-1^ (b) of 350 wheat genotypes including check (Kharchia 65). Salinity stress was imposed for 4 weeks as mix of salts to 7 days old wheat seedlings grown under hydroponics. Green circle indicate positive correlation while yellow circle indicate negative correlation. Single asterisk (*) indicate 5% level of significance, double asterisk (**) indicate 1% level of significance and three asterisks (***) indicate <1% level of significance.

### Principal component analysis of salinity tolerance associated traits

The PCA was conducted on trait data collected from wheat landrace genotypes grown under control and 15 dSm^-1^ **(Fig. 5 and Supplementary Fig. 2)**, with **Table. 1** displaying the eigenvalues more significant than one for the first four components and five components in the control group and 15 dSm^-1^ group, along by their respective variance percentages and cumulative variance percentages. In control, the total number of variables examined in this study is 18, equivalent to the number of components. In control, the biplot represents the distribution of variance across the first two principal components (PC), as well as the correlation across attributes using eigenvectors **(Fig. 5 (a))**. The first and second PCs have variances of 44.2% and 13.4%, respectively. An eigenvector indicates a correlation among the measured traits **(Supplementary Fig. 2 (a))**. A highly positive correlation was observed between SL, RL, PH, RDW, TDW, Fv/Fm, GLA, CCI, and Fv/Fm and measured MSI and WD stress-associated parameters. However, RWC correlates highly negatively with all other observed traits. The scree plot illustrates the percentage of variation across several principal components in control **(Supplementary Fig. 2 (b))**. PC1 shows a 3.29-fold higher variance compared to PC2. In 15 dSm^-1^, the total number of variables examined is 18, equivalent to the number of components. The biplot represents the distribution of variance across the first two principal components (PC), as well as the correlation across attributes using eigenvectors in 15 dSm^-1^ **(Fig. 5 (b))**. The first and second PCs have 44 % and 13.8 % variances, respectively. An eigenvector indicates a correlation among the measured traits **(Supplementary Fig. 2 (c))**. A highly positive correlation was observed between TDW, TFW, SL, RL, PH, RDW, SDW, GLA, Fv/Fm, Fv/Fm, and CCI and measured MSI, WD stress-associated parameters, but RWC and VS show highly negative correlation with all other observed traits. The scree plot illustrates the percentage of variation across several principal components in control **(Supplementary Fig. 2 (d))**. PC1 shows a 3.18-fold higher variance compared to PC2.

**Fig 5.**
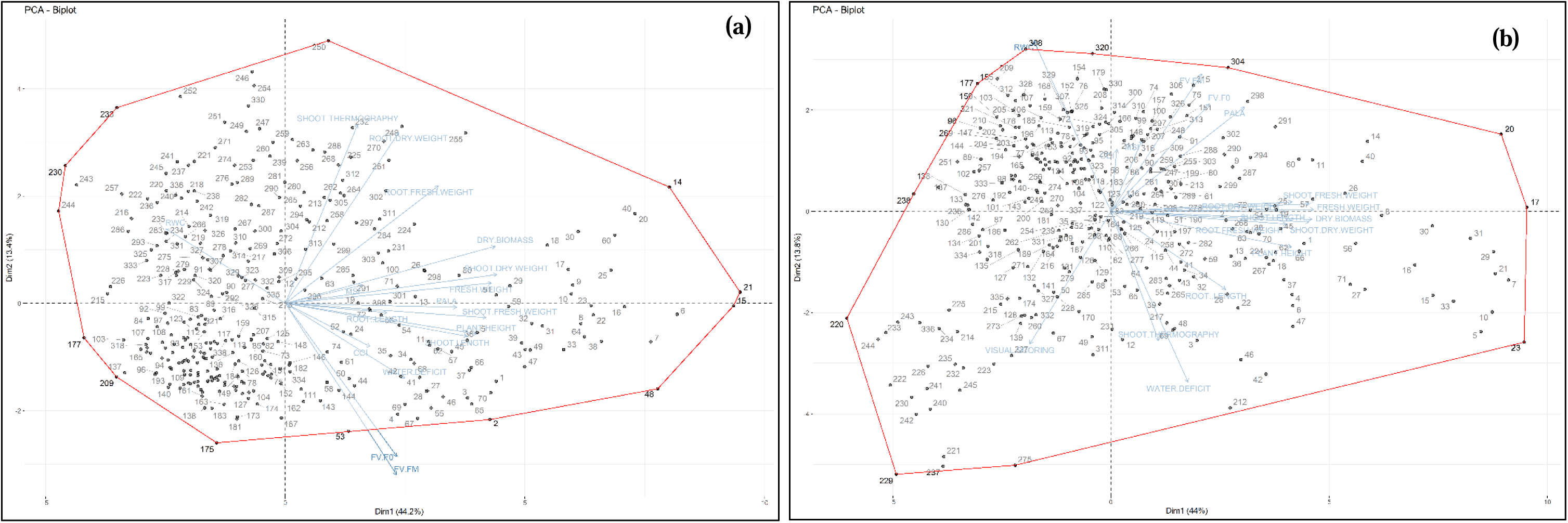
Principal component analysis of manual traits. The Biplot shows PCA of 350 wheat genotypes including check (Kharchia 65) under Control (a) and 15 dSm^-1^ (b). Salinity stress was imposed for 4 weeks as mix of salts to 7 days old wheat seedlings grown under hydroponics.

### Identification of tolerant landraces using MGIDI (Multi-trait genotype-ideotype distance index)

MGIDI index is calculated, and based on the MGIDI value, the tolerant and least tolerant genotypes are selected. The selection pressure, which accounts for approximately 15 % of the decision-making process, led to the identification of the top fifty and bottom fifty genotypes for each scenario. In control, the tolerant genotypes **(Supplementary** Fig. 3) were selected; in 15 dSm^-1^, the tolerant genotypes (**Supplementary** Fig. 4) were deemed the most effective. For tolerant genotype selection, the genotype with the lowest MGIDI score is regarded as nearer to the desired ideotype and will probably have desirable values for all the examined features. The top performing 50 genotypes have been selected based on the MGIDI score. In control, for measured traits, the genotype IC138337 has the lowest MGIDI value (1.9962), while IC397820 (4.4767) has the highest value **(Supplementary Table. 2)**. In the 15 dSm^-1^, for measured traits, the lowest MGIDI values are observed in the genotype K-65 (3.3284), whereas EC637483 (5.9633) showed the highest MGIDI values **(Supplementary Table. 3).** The least tolerant genotype has the highest MGIDI score, farthest from the desired ideotype. The lowest/poor-performing 50 genotypes have been selected based on the MGIDI score. In control, for measured traits, the selected genotypes MGIDI value ranges from 6.2800 (IC138379) to 7.8974 (IC589278**) (Supplementary Table. 4)**. In the 15 dSm^-1^, for measured traits, the selected genotype MGIDI value ranges from 8.0239 (IC26729) to 10.0460 (IC551375) **(Supplementary Table. 5).**

### The strength and weakness view of the top performing 50 tolerant genotypes selected based on the MGIDI index

The radar plot shows a visual representation of the strengths and weaknesses of the top performing 50 selected tolerant genotypes in control (**Fig. 6)** and 15 dSm^-1^ (**Fig. 7).** The strengths and weaknesses evaluate the impact of each factor on the MGIDI index in selected genotypes. This will use a ranking system to visually represent each element’s relative worth. In this depiction, variables exerting a greater influence are positioned closer to the plot’s centre, while those with lesser influence are situated towards the periphery. FA1 consists of several traits in control, including CCI, GLA, SL, PH, SFW, TFW, SDW, TDW, Fv/Fm, and Fv/F0. FA2 consisted of traits like RFW, RDW, and CT. FA3 comprised traits RWC and WD, and FA4 consisted of MSI and RL (**Fig. 6)** and **Table. 2.** The order of factors close to the centre to the periphery is FA1>FA2>FA4>FA3. FA1 is closer to the centre in the EC6903 genotype, followed by IC345620, IC138553, and FA1 of genotype IC421880 is plotted toward the peripheral, among other genotypes. FA3 plotted the peripheral region of the circle. In FA3, the genotype IC449213 is plotted extremely on the outer side of the circle, among other genotypes. In 15 dSm^-1^, FA1 consists of traits including GLA, SL, PH, SFW, RFW, SDW, and RDW. FA2 consisted of traits like RWC and WD. FA3 comprised CT, Fv/Fm, Fv/F0, FA4 consisted of VS and RL, and FA5 has CCI and MSI with positive loading (**Fig. 7)** and **Table. 3.** The order of factors close to the centre to the periphery is FA5>FA1>FA4>FA2>FA3. FA5 is closer to the centre in the IC532414 genotype, followed by IC532495, IC532099, and genotype EC637463, FA5 of genotype “1531” is plotted toward peripheral among other genotypes. FA3 plotted the peripheral region of the circle. In FA3, the genotype IC252899 is plotted extremely on the outer side of the circle, among other genotypes.

**Fig 6.**
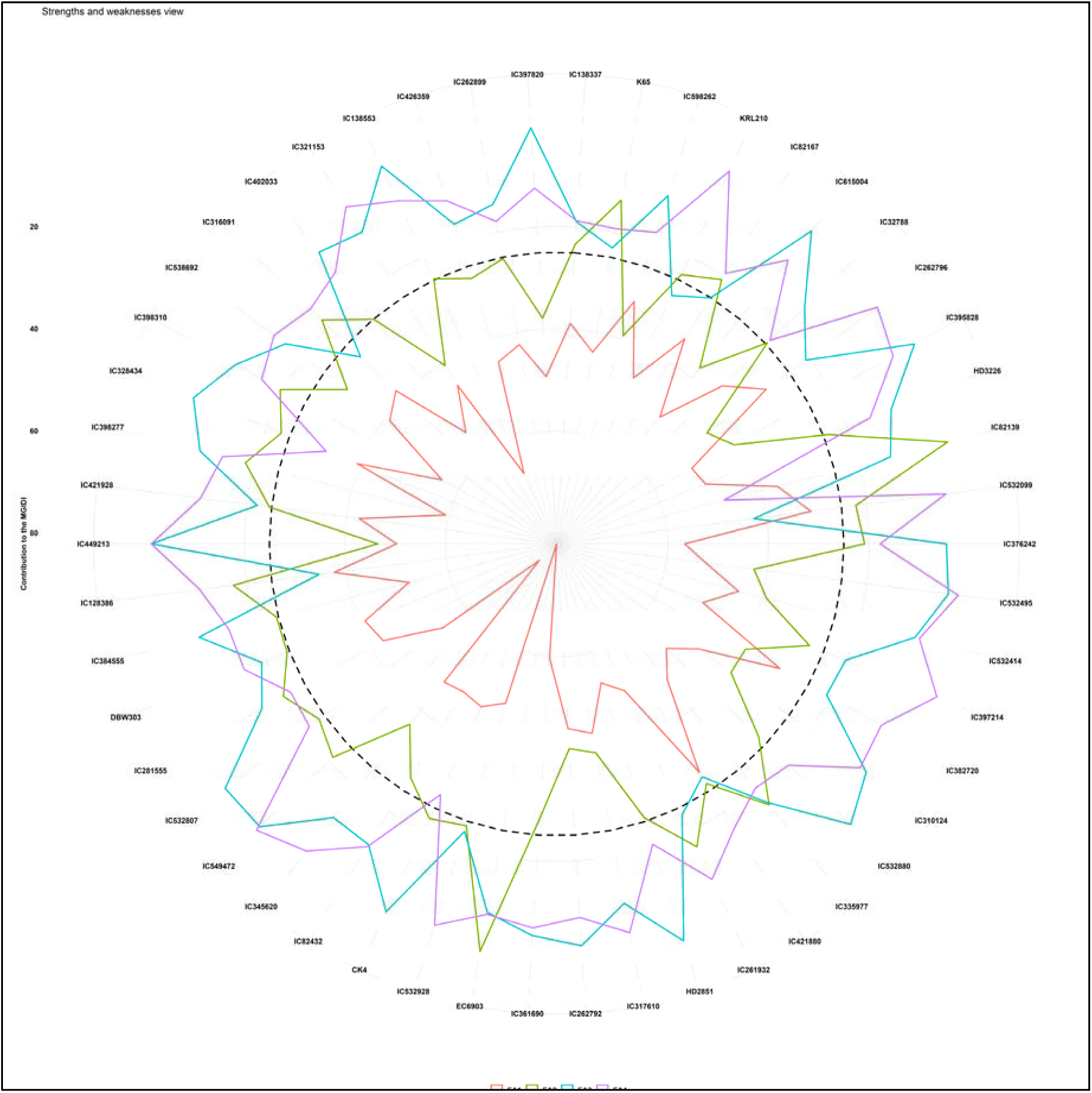
The radar plot represents the strengths and weaknesses view of the selected genotypes under control condition grown under hydroponics.

**Fig 7.**
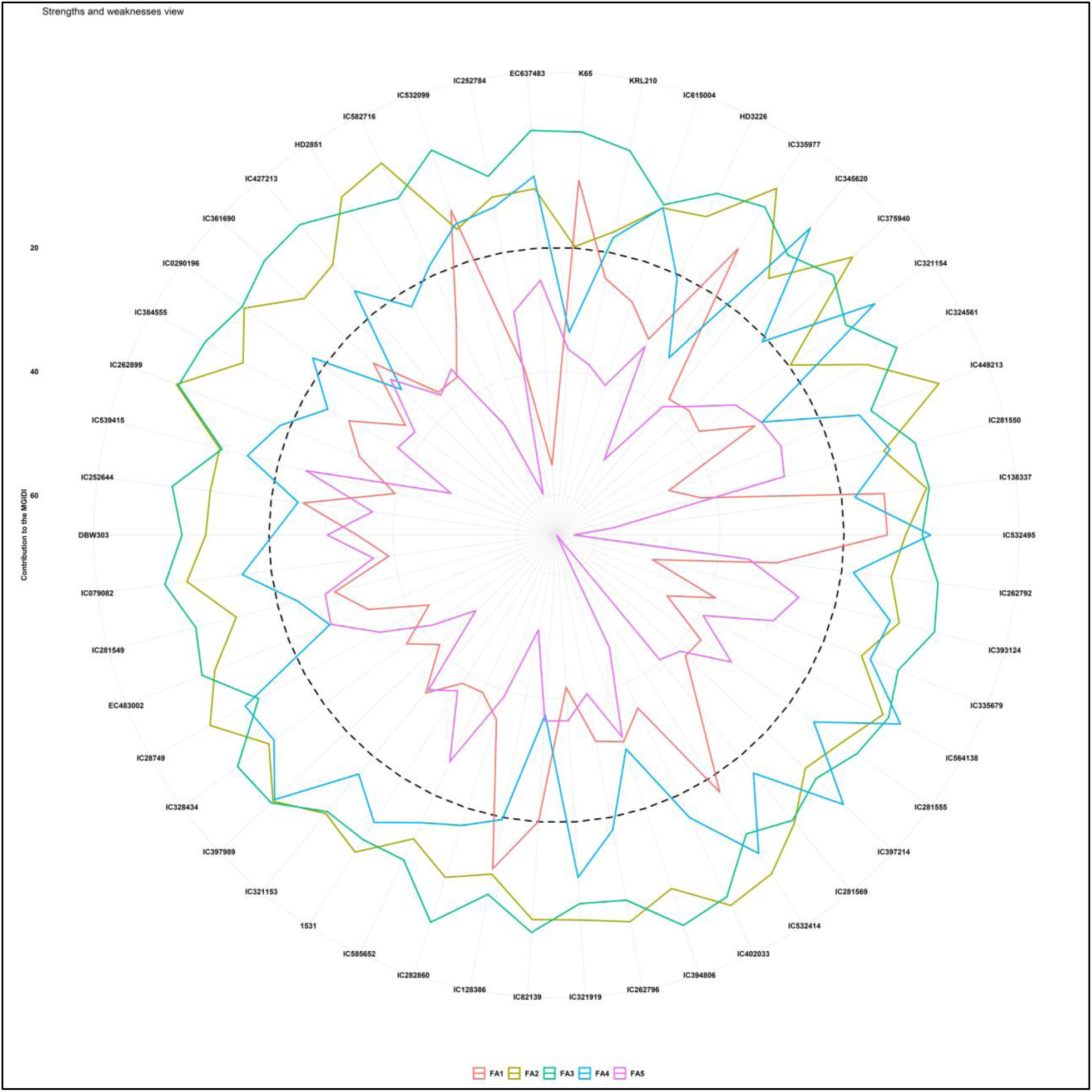
The radar plot represents the strengths and weaknesses view of the selected genotypes under salt treatment (15 dSm^-1^) grown under hydroponics.

### The proportion of phenotypic variance based on the MGIDI index

The proportion of phenotypic variance in the measured traits under control and 15 dSm^-1^ of the top performing 50 selected tolerant genotypes based on the MGIDI index (**Supplementary Fig. 5 (a, b)).** Variations were seen in the overall performance and variance components among each trait group. Furthermore, it has been shown that genetic variables play a substantial role in accounting for the variability seen in most phenotypes. In the control condition, in this single-environment study, it was shown that the genotypic component of the variance for phenological features was more significant than the residual components among all the traits, and it is below or equal to the threshold level. Among the studied characteristics, CT has a more significant genotypic variance and a lower residual variance compared to increasing order WD, RWC, GLA, and CCI, which have a higher residual variance of 0.5 with an equal amount of genetic variance (**Supplementary Fig. 5 (a**)). In 15 dSm^-1^, a single-environment study, it was shown that the genotypic component of the variance for phenological features was more significant than the residual components among all the traits, and it is below the threshold level except CCI. Among the studied traits, CT has a larger genotypic variance and a lower residual variance compared to increasing order Fv/Fm, VS, WD, RWC, MSI, others, and CCI, which have a higher residual variance of above 0.5 (**Supplementary Fig. 5 (b**)). Findings suggest that the genotypes consistently influenced the manifestation of characteristics.

### Identification of tolerant landraces using Smith-Hazel Index (SHI)

Smith Hazel index (SHI) is calculated and based on genetic worth; the tolerant and least tolerant genotypes were selected. The selection pressure, which accounts for approximately 15 % of the decision-making process, led to the identification of the top fifty and least fifty performing genotypes for each scenario. In control and 15 dSm^-1^, the tolerant genotypes were selected (**Supplementary** Fig. 6 and 7) and were deemed the most effective. The corresponding selection gain of measured traits in control and 15 dSm^-1^ are in **Supplementary Table. 6** and **Supplementary Table. 7.** For tolerant genotype selection, the genotype having the highest genetic worth is regarded as tolerant of examined features. The top performing 50 genotypes have been selected based on their genetic worth. In control, the genetic worth of selected genotypes measured traits ranges from 510.59 (IC138337) to 4.4767 (IC281555) **(Supplementary Table. 8)**. In 15 dSm^-1^, the genetic worth of selected genotypes measured traits ranges from 6953.231 (K-65) to 6888.616 (IC418384) (**Supplementary Table. 9).** For least tolerant genotype selection, the genotype with low genetic worth is regarded as least tolerant of examined features. The least performing 50 genotypes have been selected based on their genetic worth. In control, the genetic worth of selected genotypes measured traits ranges from 322.0793 (IC335684) to 280.2355 (IC111914) **(Supplementary Table. 10)**. In 15 dSm^-1^, the genetic worth of selected genotypes measured traits ranges from 6830.478 (IC536474) to 6800.458 (4379) **(Supplementary Table. 11).**

### Identification of tolerant landraces using Salt Susceptibility Index (SSI)

The stress Susceptibility Index (SSI) was calculated for each trait of all the genotypes. A total number of genotypes arranged into an ascending order based on their SSI value and rank has been given (**Supplementary** Fig. 8). In SSI, the genotype ranked first is considered less vulnerable than the genotype ranked last. In genotype, the selection process involves the identification of the top-ranked 50 genotypes exhibiting high levels of tolerance **Supplementary Table. 12**, as well as the selection of the last-ranked 50 genotypes considered least tolerant **Supplementary Table. 13**.

### Most tolerant and least tolerant landraces selected based on three indices

To identify the tolerant and least tolerant genotypes grown in salt stress conditions, the selected top-performing 50 genotypes and least-performing 50 genotypes from each indices MGIDI, SHI, and SSI in 15 dSm^-1^ were used, and a Venn diagram was prepared. The common genotypes from the top 50 and the last 50 selected genotypes from three indices are considered tolerant and least tolerant **(Fig. 8).** The results showed that out of the top 150 genotypes from all indices, 7 were common and considered tolerant **(Fig. 8 (a))** and **Table. 4** and of the last 150 genotypes from all indices 12 were common and considered least tolerant **(Fig. 8 (b))** and **Table. 4.**

**Fig 8.**
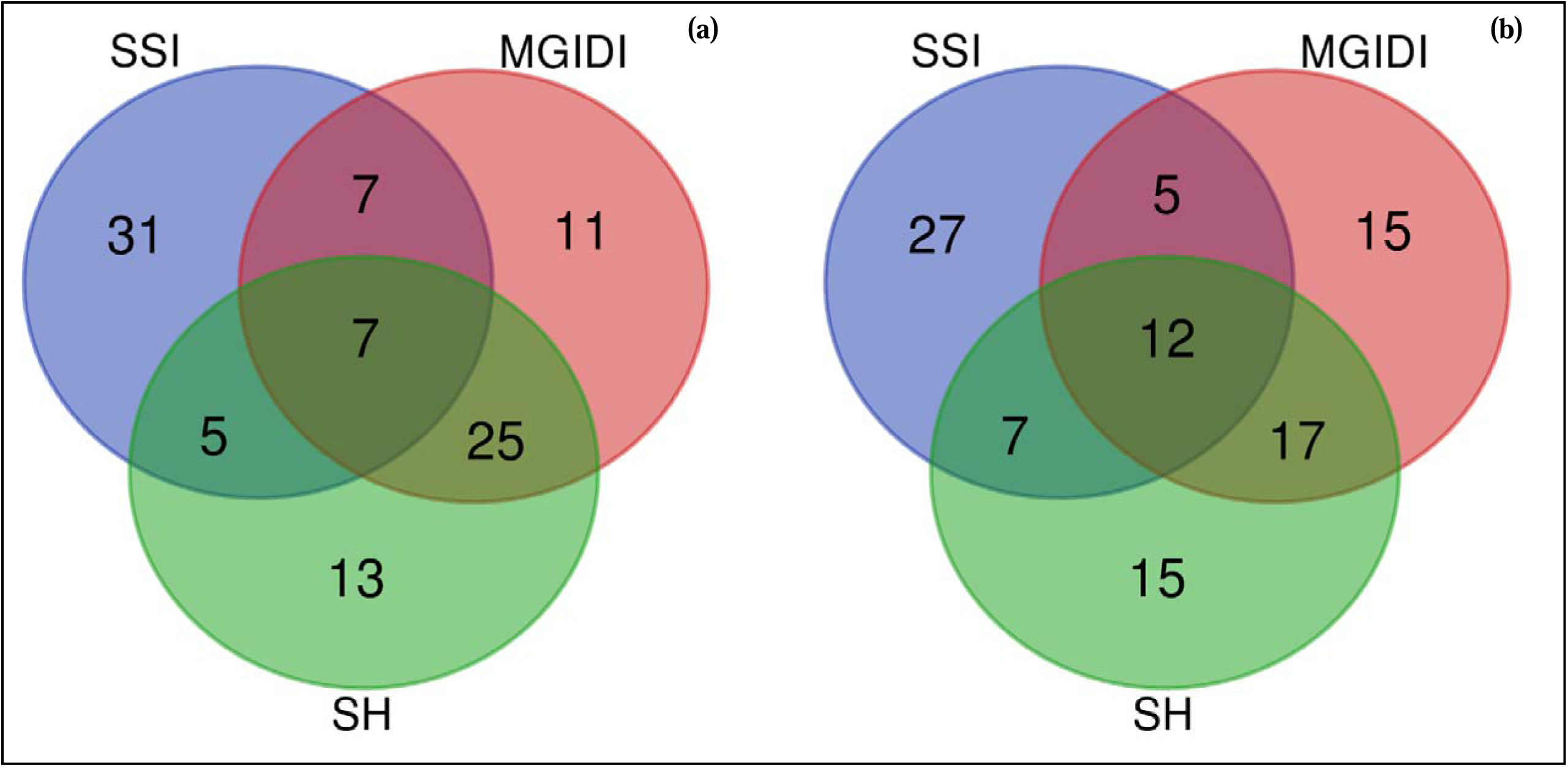
The venn diagram shows common genotypes in all three indices under salt treatment 15 dSm^-1^ of 150 wheat genotypes including check (Kharchia 65) selected based on ascending order ranking in MGIDI, SHI, and SSI. Tolerant genotypes (a), and least tolerant genotypes (b).

## 4. Discussion

Landraces are characterized by their diverse and localized adaptations of domesticated species, making them valuable genetic resources for addressing the demands of agriculture in challenging areas. The local ecotypes exhibit phenotypic variability and possess a moderate edible yield while concurrently demonstrating good nutritional value. The primary contributions of landraces to the field of plant breeding encompass the provision of features that enhance nutrient absorption and utilization efficiency and the provision of valuable genes that facilitate adaptation to challenging environmental conditions, including water stress, salt, and high temperatures (Dwivedi *et al*., 2016). The landraces of wheat, which have considerable potential as a source of genetic diversity for enhancing the crop, have received less attention in investigating individual features (Ehdaie and Waines, 1989; Munns and James, 2003). The morphological and stress-associated traits were recorded in 3 weeks of wheat landraces grown under the treatments control and 15 dSm^-1^. The rapid maturation of wheat caused by saline stress decreased crop height and leaf area (Ahmad *et al*., 2013). The length of the plumule was observed to be highly susceptible to salt stress (Bacilio *et al*., 2004).

Excess salt stress in the early vegetative phase significantly impacts wheat growth, including root patterning, leaf expansion, and leaf area. In addition, the study conducted by (El-Hendawy *et al*., 2009; Borlu *et al*., 2018) revealed that salt stress had a detrimental effect on the number of leaves per plant, leaf size, and leaf lifetime, causing them to shrink. A decrease in the growth of plants was noticed in wheat landraces subjected to 15 dSm^-1^. In accordance with previous investigations, all evaluated wheat accessions in the present study exhibited a significant decrease in plant growth compared to the control plants. A variation in genotypes was identified regarding growth response at 15 EC in comparison to tolerant and sensitive checks. The shoot length, root length, and plant height were measured under high salt stress treatment, and the statistical significance of these measurements were also determined. This finding suggests that the seedling establishment stage is particularly vulnerable to salt stress among different phenophases of wheat. Under stress conditions, seed germination and the early stages of seedling development are the most crucial for plant survival. El-Hendawy et al., 2017 and Turki *et al*., 2012 found positive associations between salinity tolerance parameters recorded during the early growth stages and those measured at full maturity. In addition, (Dugasa *et al*., 2016) and (Hussain *et al*., 2021) revealed a correlation between morphological parameters evaluated in the tillering phenophase and a reduction in wheat grain yield (phenophase of full maturity). Among the various accessions studied, only a few demonstrated satisfactory growth under salt-stressed conditions, indicating the presence of genetic traits that confer potential tolerance to initial salt stress in wheat landraces. The bread wheat genotypes exhibited substantial variation in root and shoot biomasses (RDW and SDW). Under salt stress conditions, the RDW SDW of the 20 evaluated genotypes exhibited a drop of 24.44% and 49.98%, respectively (Pour-Aboughadareh *et al*., 2021). Our findings revealed that the genetic background of the genotypes under investigation significantly influenced the impact of salt stress on the response of wheat plants. This was evident via the significant variations identified for SFW (57%), RFW (48%), TFW (54%), SDW (44%), RDW (50%), and TDW (47%) in studied genotypes in 15EC.

The Chlorophyll Content Index (CCI) was utilized to quantify the relative value of plant chlorophyll (Fanizza *et al*., 1991). The CCI exhibited a negative association with the intensity of salt stress across all wheat genotypes (Saddiq *et al*., 2021). Our findings indicate a significant drop was observed in studied landraces in salt-stressed conditions compared to the normal-grown plants. The imposition of salt stress resulted in a notable reduction in leaf area compared to the control plants. However, it is essential to note that genotype differences were noted (Khan *et al*., 2010). We also observed similar results in our studied landraces, and the reduction in GLA ranged from 0.00 cm^2^ of 27 genotypes to 44.25 cm^2^ (K-65) with an average mean of 12.82 cm^2^. The findings suggest that the reduced biomass accumulation in salt-treated plants can be attributed to a decrease in chlorophyll levels and a decrease in green leaf area. The increasing level of the salt concentration in the soil solution leads to early osmotic stress, which subsequently hinders the plant’s ability to absorb water for its transpiration requirements (Seyed Sharifi *et al*., 2017). In order to ensure optimal metabolic processes, plants must effectively manage cellular water levels to cope with stress. In salt stress conditions, the genotype K-65, known to be salt tolerant, exhibits a higher relative water content (RWC) than the sensitive genotype HD2687 (Sairam *et al*., 2005). The results of our study indicate that accession IC418402 had the highest RWC of 91.49% under salt stress conditions, demonstrating its ability to maintain a minimal water deficit. This implies that particular genotypes can maintain cell turgidity to ensure optimal functionality in stressful conditions. During stress, the cell membrane must uphold its structural integrity while preserving its inherent properties to ensure optimal functionality (Khatkar and Kuhad, 2000). A higher concentration of hydrogen peroxide (H_2_O_2_) under stress triggers a cascade of lipid peroxidation events, resulting in detrimental effects on the cellular organelle membrane. Ultimately, this leads to the loss of cellular functionality (Singh *et al*., 2020). The findings of our study indicate that elevated salt stress conditions result in membrane impairment. Specifically, our results demonstrate that plants subjected to a salinity level of 15 dSm^-1^ exhibited more significant electrolyte leakage than plants not exposed to salt treatment. The accession IC7145972 exhibited a relatively low level of electrolyte leakage, measuring at 5.72% when exposed to a salinity level of 15 dSm^-1^ for an extended amount of time. This implies the ability to effectively preserve the integrity of the cell membrane in challenging environmental conditions.

Visual salt injury was recorded based on visual scoring in rice genotypes grown in salt treatment (Suriya-arunroj *et al*., 2004). The score ranges from 1-9 and is given based on the necrotic level of leaves under salt-stressed conditions and the extent of necrotic regions: Score 1 denoted the absence of necrotic areas, score 3 indicated necrotic areas ranging from 1% to 25%, Score 5 represented necrotic areas ranging from 26% to 50%, score 7 encompassed necrotic areas ranging from 51% to 75%, and score 9 indicated necrotic areas amounting to 76% to 100% or plant death (Zhen *et al*., 2010). Our findings also support the observation that very few landrace accessions consistently exhibited reduced leaf necrosis (50%-70%) and improved growth when subjected to salt stress conditions. The study intended to assess the genetic variability in the response of seedling leaf temperature to osmotic stress caused by a concentration of 150 mM NaCl in 18 different wheat genotypes. The primary method of measurement used in this evaluation was IR thermography (Sirault *et al*., 2009). Our results show landrace accessions showing higher the canopy temperature than controlled plants. The inability of seedlings to extract water from a salt solution during the early stage of osmotic stress can be attributed to the water potential difference between the root cortex and the salt solution (Said *et al*., 2022). Consequently, plants will close their stomata, resulting in a decrease in evaporative cooling and, ultimately, an increase in CT. The quantification of salt-induced damage in the photosynthetic system may be effectively assessed by measuring chlorophyll fluorescence (Baker and Rosenqvist, 2004). ROS damage the thylakoid membrane of PSII under stressful salt condition. Our data showed that the landraces showed a decrease in Fv/Fm compared to the control plants, with a value of 0.8 in the control plants.

The correlation pattern in this study also showed a positive association between TDW, RL, SL, RFW, SFW, TFW, RDW, SDW, CT and WD under both control and salt stress conditions. Remarkably, dry biomass emerged as a prominent factor that had a strong positive association with most other growth metrics linked to biomass under both salt and control conditions **(Fig. 4).** Principal Component Analysis (PCA) was employed in our research to ascertain the primary selection qualities that contribute significantly to salt tolerance. The first and second principal components were utilized for this purpose **(Fig. 5 and Supplementary** Fig. 2(b) and 2(c)). PCA-biplot is a multivariate analytic technique that integrates features and objects into a two-dimensional representation, thereby reducing overlapping variations. PCA thus facilitates the identification of critical characteristics for selection (Arzu *et al*., 2018). The characteristics SDW, TDW, TFW, RFW, SFW, RL, PH, SL, CT, MSI, and Fv/Fm were shown to contribute more to characterizing the variance between cultivars, according to the PCA in control and salt stress conditions.

In contrast, the observed augmentation in the overall dry mass may be attributed to the combined growth in RL, SL, RFW, SFW, RDW, and SDW. The combined findings of PCA and correlation analysis suggest that certain morphological and stress-associated features, such as TFW, TDW, MSI and CT, can serve as valuable indicators in identifying and selecting better genotypes with enhanced salt tolerance. These findings are supported by (Al-Ashkar *et al*., 2020; Maqbool *et al*., 2020).

### Donor selection based on genetic indices

Salinity is related to the level of mineral salts that are dissolved in the soil solution. Soils are categorized under saline stress when they hold a concentration of salt that is significant enough to hinder the growth of most crop species. Saline soils can be characterized by their electrical conductivity values surpassing 4 dSm^−1^ (Shahzad *et al*., 2019). A rise in soil salinity induces three separate forms of stress in plants: osmotic, ionic, and oxidative (Pan *et al*., 2020). The plant’s ability to uptake water from the soil solution is reduced due to osmotic stress, resulting in a water deficit. Conversely, ionic stress is attributed to an increased concentration of Na^+^ and Cl^-^ ions inside cellular components involved in metabolic processes, impacting enzymes’ functioning. Excessive Na^+^ in the soil can cause nutritional shortages, namely in K^+^, by impeding the absorption of potassium and promoting its exclusion from cellular structures (Ma *et al*., 2016).

The initial impact of salinity stress on plants is osmotic stress, followed by ionic and oxidative stress (Bagwasi *et al*., 2020). Consequently, plants respond by closing their stomata, reducing transpiration rate, and experiencing a loss of turgor pressure. This interruption in turgor pressure hinders roots’ expansion and shoot cell growth (Zhao *et al*., 2020). During the subsequent phase of growth reduction, salts accumulate within plant leaves, eventually reaching critical levels that lead to toxicity (Al-shareef and Tester, 2019). Excessive salt concentration beyond an optimal threshold harms wheat plant growth, leading to various adverse effects such as diminished leaf area, decreased total biomass, shorter shoot and root lengths, and a decline in the relative growth rate (Yang *et al*., 2009). Salinity stress leads to the down-regulation of various genes responsible for developing branch and spikelet meristems, reducing the number of grains (Zheng *et al*., 2021). In saline conditions, various plant yield components, including seed production, spike length, spikelets per spike, and 1000 grain weight, were significantly and adversely impacted (Mishra *et al*., 2014). Under saline irrigation, the allocation of plant dry matter was disrupted, leading to a decrease in plant height, leaf area, and the dry weight of both shoots and roots (Abou-Baker and El-Dardiry, 2015). The harvest index exhibits variability ranging from 0.2 to 0.5 in wheat (Husain *et al*., 2003). A low salt level may not significantly impact grain production despite reducing leaf area and shoot biomass. It results in a rise in harvest index with salinity and a finding that grain yield does not decline. A significant amount of genetic diversity was seen in the barley and durum wheat crops cultivated under drip irrigation conditions with varying degrees of salt, reaching up to 26 dSm-1(Royo and Abió, 2003). The data gathered at CIMMYT indicates that around 8-10% of the cultivated land dedicated to wheat production in India, Pakistan, Iran, Egypt, Libya, and Mexico experiences the detrimental effects of salt (Mujeeb-Kazi and De Leon, 2002). Wild relatives, such as some halophytes, have the potential to provide valuable sources of tolerance for enhancing wheat production. A unique multi-trait genotype-ideotype distance index (MGIDI) aims to facilitate genotype selection and treatment recommendation using information from several variables. The performance of the proposed index is tested using Monte Carlo simulations. These simulations calculate the success rate of picking characteristics with desired gains under different situations. The scenarios include variations in the number of genotypes, evaluated traits, and the correlation structure between traits (Olivoto and Nardino, 2021). A study assessed the multi-trait stability index (MTSI) of 45 maize inbred lines. The MTSI was derived from the examination of 18 distinct attributes. Nine inbred lines were chosen using a selection process characterized by a selection intensity of 20% (Balbaa *et al*., 2022). Using Multivariate analysis in Recombinant Inbreed Lines (RIL) for stay green trait and stem reserve mobilization under drought and heat stress (Taria *et al.,* 2023) The study based on MGIDI showed that a total of eight genotypes in Season 1 and Season 2, as well as three genotypes in Season 3, displayed favourable performances and were selected for QTL mapping to Phenology-Affecting Genes (Vrn, Ppd, and Eps) in bread wheat (Farhad *et al*., 2023). The genotypes with the highest and lowest rankings were chosen by employing the MGIDI score, which considers many factors seen in wheat seedlings cultivated under hydroponics. The top fifty and least fifty genotypes were considered tolerant and least tolerant, respectively grown in 15 dSm^-1^ (**Supplementary** Fig. 3 and 4**, Table. 3 and 5**). (Smith, 1936) proposed a linear index of weighted phenotypic characteristics for wheat line selection. (Hazel, 1943) introduced a method for determining genetic variances and covariances, which helped determine index weights. The index weights optimize the correlation between phenotypic qualities and aggregate genotype, which is a weighted collection of unknown genetic values based on economic value. Selecting using a Smith-Hazel index maximizes genotype gain when genetic variances and covariances are accurately estimated. Enhancing the Genetic Composition of a Breeding Population of Spring Wheat via SHI (Wells and Kofoid, 1986). The genetic response to SH index selection for grain production, kernel weight, and percentage of protein in four wheat hybrids (Gebre-Mariam and Larter, 1996). The study evaluated the yield and quality characteristics of sweet corn, and the selection of the top performance was determined using the Sweet Corn SH index (Asghar *et al*., 2010). In our study, the top 50 and bottom 50 wheat landraces were selected based on their genetic worth, as determined by the SHI method (**Supplementary** Fig. 6 and 7). The top 50 landraces were considered tolerant genotypes, while the bottom 50 were considered the least tolerant genotypes grown in 15 dSm^-1^ (**Supplementary Table. 9 and 11)**.

Multiple stress-associated features were utilized to select wheat genotypes with salt tolerance, which were then cultivated under salinity conditions (Singh *et al*., 2015). SSI was used to choose wheat genotypes that performed best when grown in salt-stress conditions (Goudarzi and Pakniyat, 2008). The high-yielding wheat genotypes were selected from the population cultivated under salt stress conditions using SSI (Hasan *et al*., 2015). Here in our study, we selected the top 50 SSI-ranked landraces regarded as tolerant genotypes, while the bottom 50 SSI-ranked were considered the least tolerant genotypes ((**Supplementary** Fig. 8**) and (Supplementary Table. 12** and **13))**.

Despite being moderately salt tolerant, salinity levels beyond 10EC significantly reduces wheat yields (Munns et al. <u>2006</u>). Globally Kharchia 65 is one of the donor line used for breeding of salt tolerant varieties. Identification of a new donor line for salt tolerance is imperative to combat the increasing problem of salinity stress. Three hundred fifty wheat landraces and check lines were evaluated for salt stress (15 dSm^-1^) tolerance in a hydroponic set-up. Across the MGIDI, SHI, and SSI, Our results showed that out of the top 150 genotypes from all indices, seven were common and considered tolerant **(Fig. 8 (a))** and **Table. 4** and of the last 150 genotypes from all indices, 12 were common and considered least tolerant **(Fig. 8 (b))** and **Table. 4.** These identified tolerant landraces serves as important resources and novel salt tolerance donors for wheat. The tolerant*sensitive, tolerant* tolerant check, crosses can be used for generating RIL populations to delineate genetic basis of salt tolerance of newly identified lines.

## Declarations

### Author Contributions

J.B. conducted the experiments and did statistical analysis and prepared the graphs. RR contributed to statistical analysis. RP contributed to measurement of physiological traits. L.S. and J.B. wrote and revised the manuscript. LS designed the experiments and acquired funding. LS, AK, SKJ, SK, ReP, AS, SuK, GPS and VC conceived the idea. All authors have read and agreed to the published version of the manuscript.

### Funding

Research was supported by the lndian Council of Agricultural Research, Department of Agricultural Research and Education, Government of lndia. This research was funded by [ICAR-IARI institute project] grant number [CRSCIARISIL20210024309] and [DBT] grant number [No.BT/Ag/Network/Wheat/2019-20].

### Data Availability Statement

All data generated or analysed during this study are included in this published article [and its supplementary information files].

## Acknowledgments

The authors thank the ICAR-Indian Agricultural Research Institute for providing the necessary facilities. JB acknowledges ICAR-IARI for the fellowship support received during the study.

## Competing interests

The authors declare that there are no competing interests

## Notes

### Competing Interest Statement

The authors have declared no competing interest.

